# Genes Encoding Xenobiotic Detoxification Proteins evolve by gene death, duplication and positive selection

**DOI:** 10.1101/2025.05.31.657154

**Authors:** Saadia Bahmmou, Sophie Fouchécourt, Philippe Monget

**Affiliations:** Physiologie de la Reproduction et des Comportements, UMR INRAE, CNRS, Université de Tours, 37380 Nouzilly, France

## Abstract

There was a huge variability in the pharmacokinetics of drugs between species, which means in the way it will be transformed, degraded and eliminated, as well as the variation in drug absorption, plasma concentration over time, half-life, bioavailability, volume of distribution, metabolism rate, and routes of excretion. To understand the reasons of such a variability between species, we have studied here the evolution of genes encoding drug-metabolizing enzymes, i.e. the 9 key genes that are known to play a principal role in this process: UGT1A6, UGT1A, CYP2B, CYP2C, CYP2D, UGT2B, CYP3A, NAT1, GSTP1. We show here that a lot of these genes have been lost during evolution in several vertebrate species: UGT1A6 in gorilla, cat dog, ruminants, pig, UGT1A in artiodactyla, UGT2B in all vertebrate species except human and gorilla, CYP2C in all vertebrate species except primates and mouse. Several of these genes have duplicated such as UGT1A in human (4 copies), CYP3A in most of vertebrate species studied here except the cat, CYP2D in the mouse (9 copies). Furthermore, several of these genes undergone evolution by positive selection such as CYP2D6, UGT1A, CYP2C (particularly in squirrel), UGT1A. Overall, this study shows that the evolution by gene death, gene duplication, and positive selection is partly responsible for the great variability in the ability of vertebrate species to metabolize drugs.

## INTRODUCTION

Very early from the 80s, scientists have consistently observed that, upon administrating drugs to different species, either orally or intravenously, there was a huge variability in the resulting pharmacokinetics between them, which means in the way it will be transformed, degraded and eliminated, as well as the variation in drug absorption, plasma concentration over time, half-life, bioavailability, volume of distribution, metabolism rate, and routes of excretion^1^, ^2^. Such variability brings up a huge challenge in the industry of pharmacology, and so, a drug tested on a specific animal on laboratory conditions cannot be reliably extrapolated to other species, as results from standard animal models, like rodents, cannot always predict other species’ responses. Considering the work and the process it takes to bring every drug to life, the pharmacology industry not only loses on terms of time and resources but also in financial terms.

To address this issue, researchers around the world still try to investigate this phenomenon from different points of view. As for this paper, we will focus on the evolutive dimension, particularly of genes that code for drug-metabolizing enzymes, to test the hypothesis that 1) several genes have been lost by pseudogenization during evolution, 2) other genes have been duplicated, and 3) their rapid and divergent evolution under the influence of positive selection is one of the causes resulting in this wide variability of activity and expression among domesticated species and humans.

For this, we have selected 9 key genes that are known to play a principal role in this process: UGT1A6, UGT1A, CYP2B, CYP2C, CYP2D, UGT2B, CYP3A, NAT1, GSTP1.

Here is a brief explanation on the roles of these Enzymes.

The cytochrome P450 (CYP) family, whose members are necessary to phase I metabolism and metabolizes over 50% of clinical drugs, involves oxidation, reduction, or hydrolysis reactions that introduce or expose polar functional groups in drug molecules, also affecting drug responses by influencing drug action, safety, bioavailability, and drug resistance through metabolism^3^.

Equally important are the phase II conjugation enzymes, such as UDP-glucuronosyltransferases (UGTs), glutathione S-transferases (GSTs), and N-acetyltransferases (NATs)^4^. These enzymes add polar groups to drugs, typically enhancing their solubility and facilitating their excretion from the body^4^.

## MATERIALS AND METHODS

### TBlastn to search for pseudogenes

The nucleic and protein sequences used for our analysis were retrieved from the Ensembl database. In search for traces of dead genes or pseudogenes, the protein sequences were blasted against the genome of domesticated species that are not present on the corresponding gene tree, using tBlastn of the BLAST programs. A figure was then constructed for each gene family, making sure that said pseudogenes, if detected, are within the conserved syntenic region, which was verified using the Ensembl browser: Genomicus.

### Codeml for positive selection

Multiple Sequence Alignments of the proteins were performed with the MUSCLE software^5^. The result files were then used to construct a phylogeny tree with the program RAxML^6^, and then to generate the nucleotide alignments with Pal2nal^7^, adding to the corresponding nucleic sequences, to generate an accurate codon alignment.

Positive selection was analyzed in both paralogs and orthologs of domesticated species that are present in the gene family trees. This included the CYP2D6 genes in cat, human, cow and mouse paralogs, the CYP2B in rabbit, human, rat, pig, goat, cat, the CYP3A in mouse paralogs, human paralogs, goat paralogs and cat paralogs, the CYP2C in rat paralogs, squirrel paralogs, mouse paralogs, human paralogs, gorilla paralogs, rabbit paralogs, the UGT1A in koala, dog, human paralogs, squirrel, mouse paralogs, bushbaby paralogs, UGT2B in human paralogs, gorilla paralogs, rabbit paralogs, squirrel, GSTP1 genes in human, gorilla, rabbit, mouse paralogs, NAT1 genes in mouse paralogs, rat, human paralogs, gorilla, pig paralogs.

Positive selection was tested using the CODEML application of the PAML packages^8^ (v 4.10.7), under the site model that allows ω (the nonsynonymous/synonymous rate ratio) to vary among codons, and the branch-site model that allows different ω values to differ among branches and among sites, with all alignment gap sites removed.

For the site model, we used 2 pairs of models: The M1a (nearly neutral) vs M2a (positive selection), and M7(beta) vs M8(beta&ω). For the branch-site model, two models were compared: the null and alternative models. In the alternative model, four site classes were defined (ω0: dN/dS<1, ω1: dN/dS=1, ω2a and ω2b: dN/dS > 1). The null model constraints ω2 = 1 to test against the hypothesis of positive selection.^9^

### Chimera to visualize the positive selected sites

Sites identified as evolving under positive selection (using BEB : Bayes empirical bayes) were mapped onto known protein structure of CYP2D6 and visualized by Chimera^10^, for better view and insight of the potential impact on protein function. The PDB database offers numerous relevant protein structures, but we selected the structure with ID **3TDA**^11^ due to its high resolution and its inclusion of the heme group, the zinc ion and the potential inhibitor prinomastat (PN0), that is in competition with thioridazine, thus providing a clearer view of the protein-ligand interactions and conformational dynamics.

A similar approach was applied to CYP2C42 of the squirrel. Its structure was retrieved from the AlphaFold Database (ID **I3LWV2**) with an average pLDDT score of 91.66.

## RESULTS

### Pseudogenes and analysis

On a global note, the gene families show a variability in the duplication events as well as pseudogenization events.

The UGT family underwent a loss of genes in most ungulates, rodents and carnivores, for instance the UGT2B gene tree, being the smallest out of the three gene families, showcases only few primates (including human and gorilla) and very few of the family of Bovidae. This loss of genes is due to a pseudogenization event, with pseudogenes being found in the conserved syntenic region, between SULT1B1(sulfotransferase family 1B member 1) and TRMPSS11E (transmembrane serine protease 11E) genes. The lack of the UGT2B paralogs is replaced by other genes, in particular by the UGT2A1 gene in the cow (fig1), UGT2A2 in goat, cat and horse, UGT2A1 and UGT2A3 in mouse.

**Fig 1:**
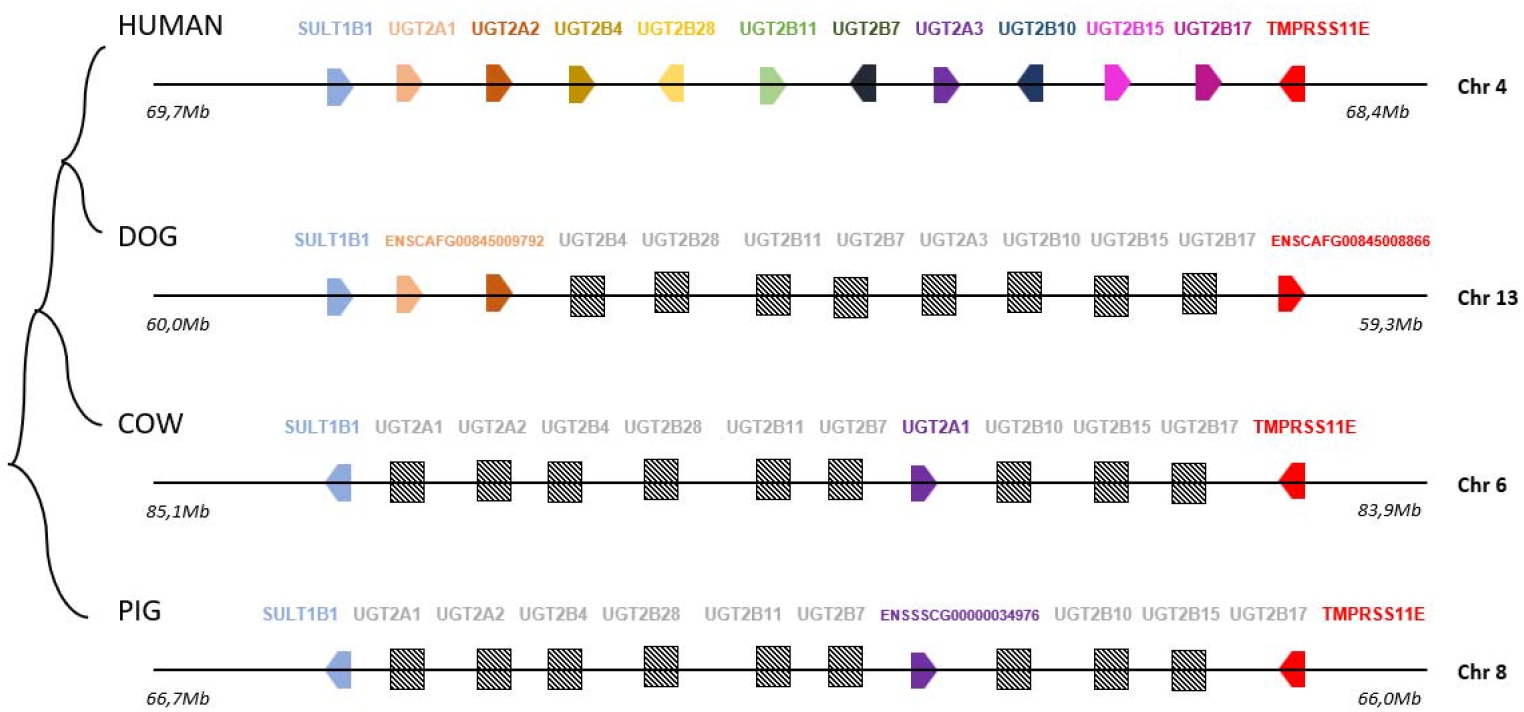
a representation of the UGT genes and pseudogenes across humans, dogs, cows and pigs, as well as their chromosomal locations : pseudogenes detected by tBlastn

**Fig 2:**
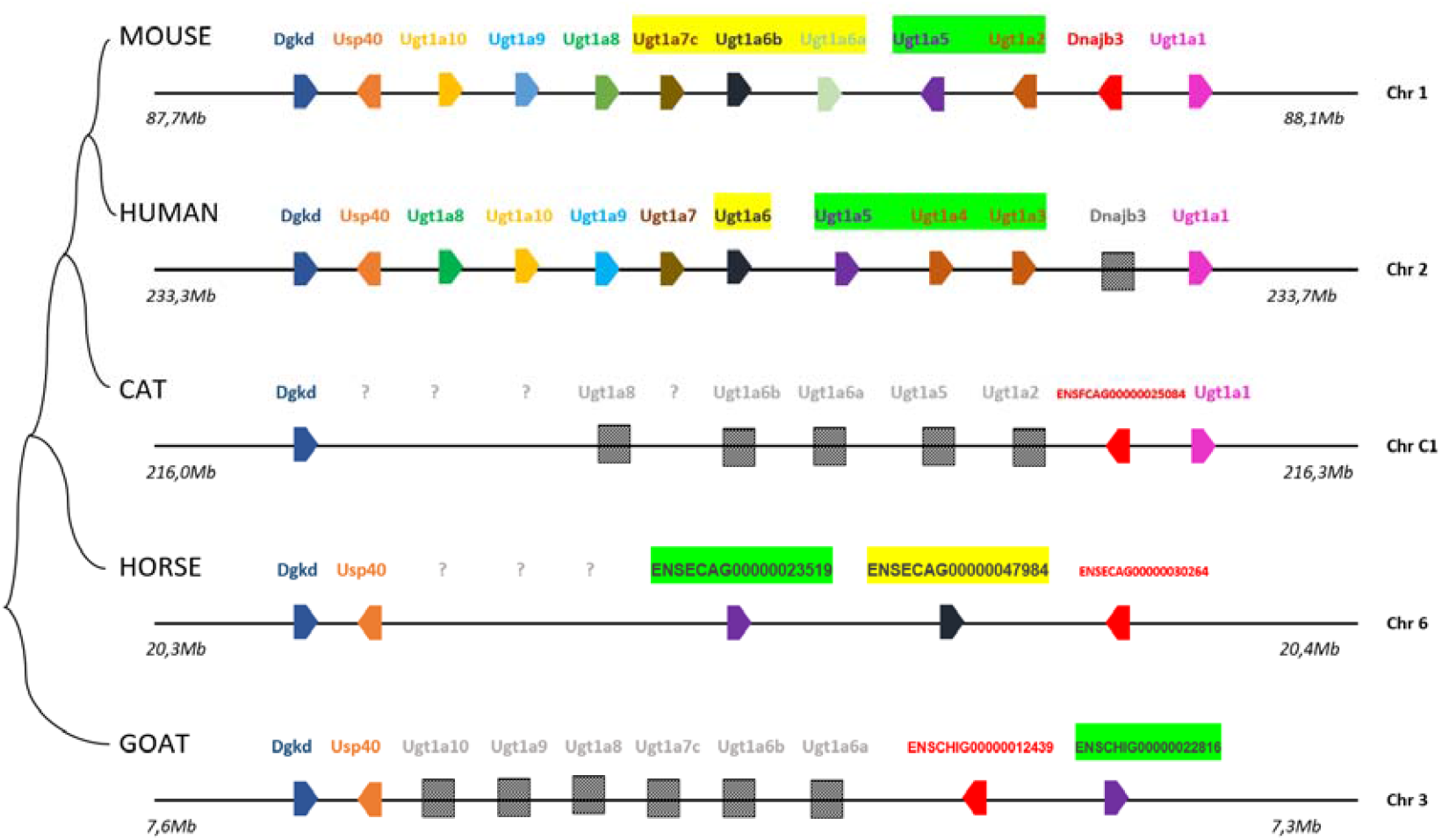
a representation of the UGT1A genes and pseudogenes across humans, mice and cats, as well as their chromosomal locations : pseudogene detected by tBlastn ?: pseudogene not detected

**Fig 3:**
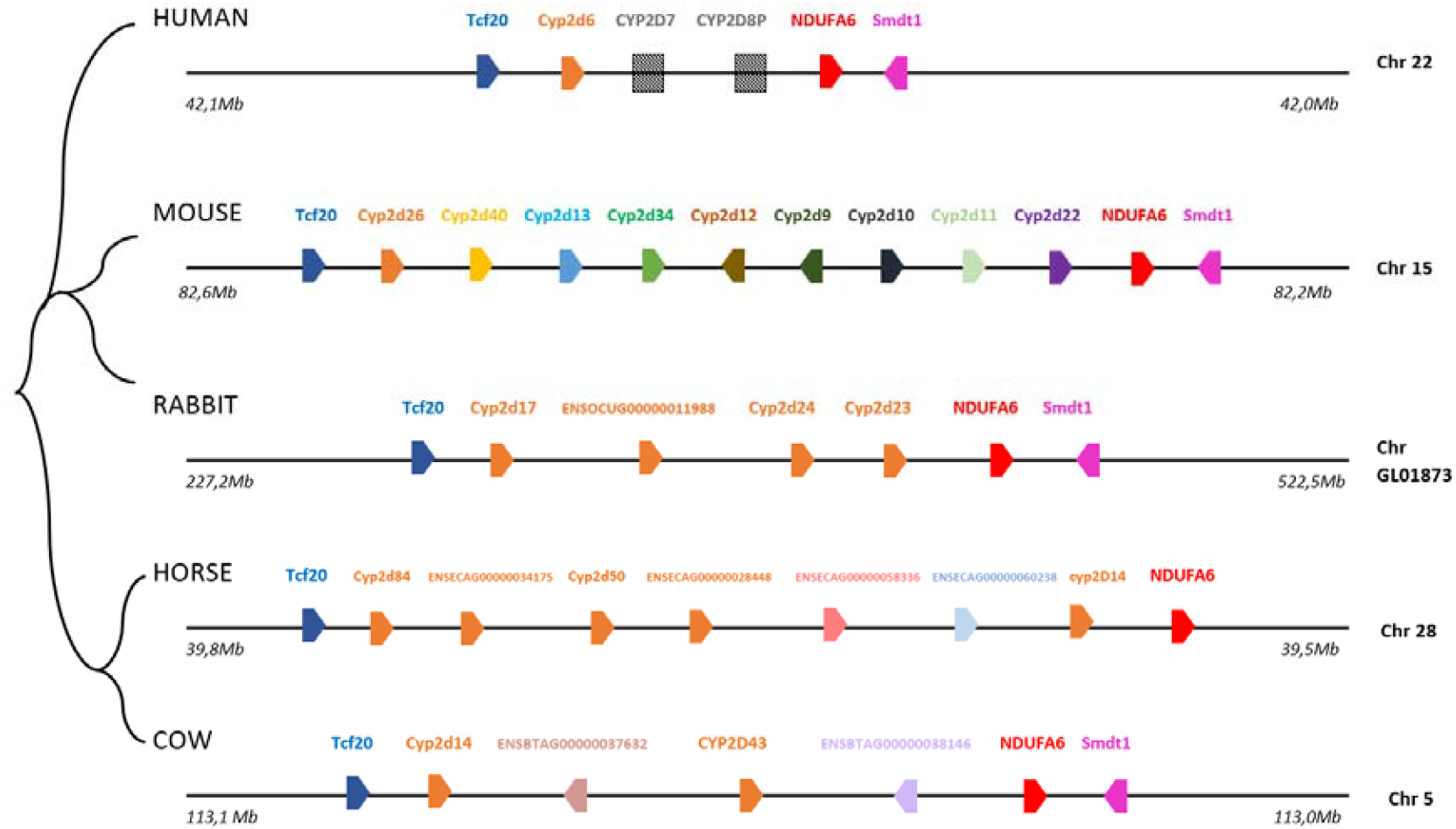
a representation of the CYP2D genes and pseudogenes across humans, mice, rabbits, horses and cows, as well as their chromosomal locations : pseudogene detected by tBlastn

**Fig 4:**
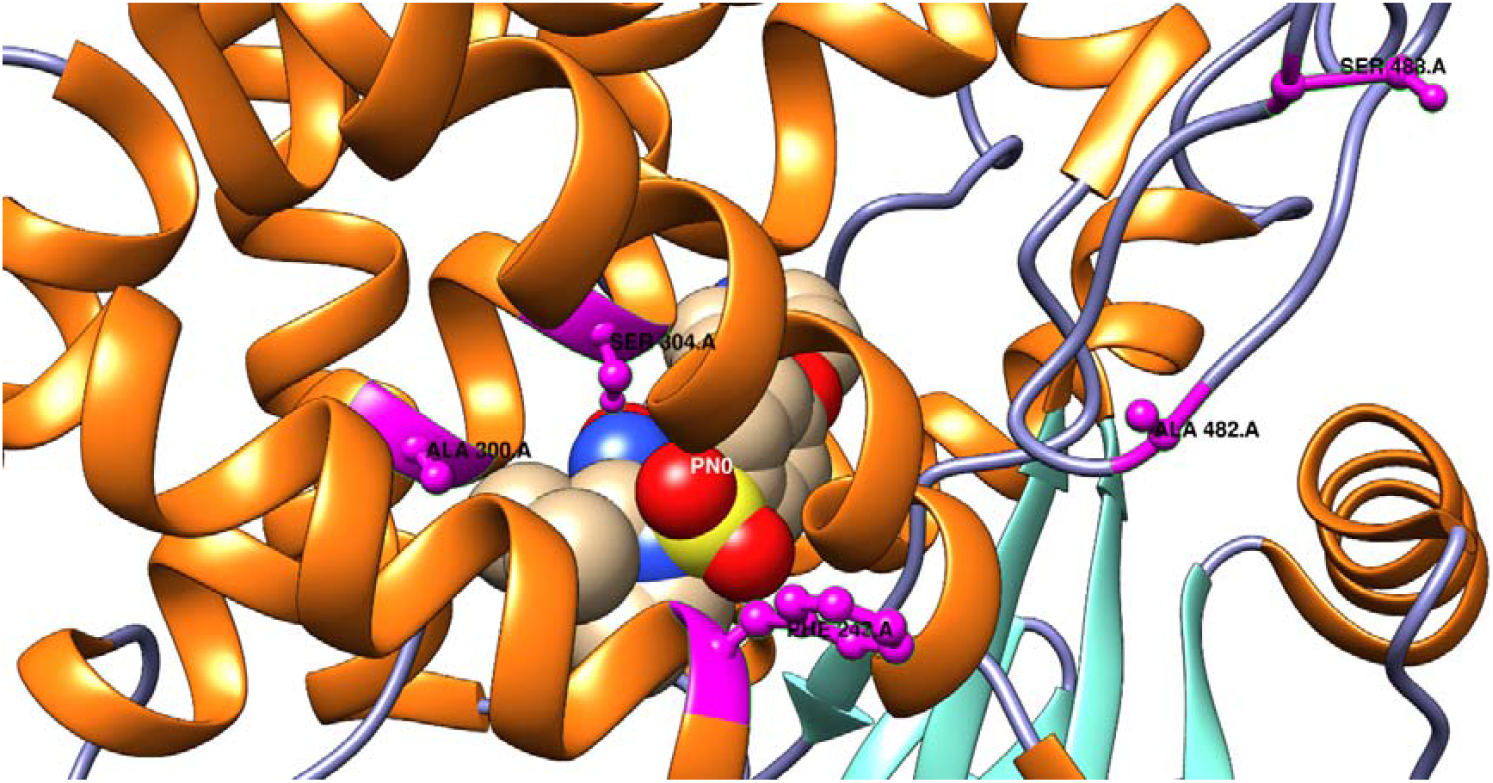
positively selected sites (BEB) represented in the 3TDA PDB structure of CYP2D6 in human The positively selected residues (81S, 300A, 304S, 138A, 142L) are colored in magenta. The inhibitor is in sphere visualization.

**Fig 5:**
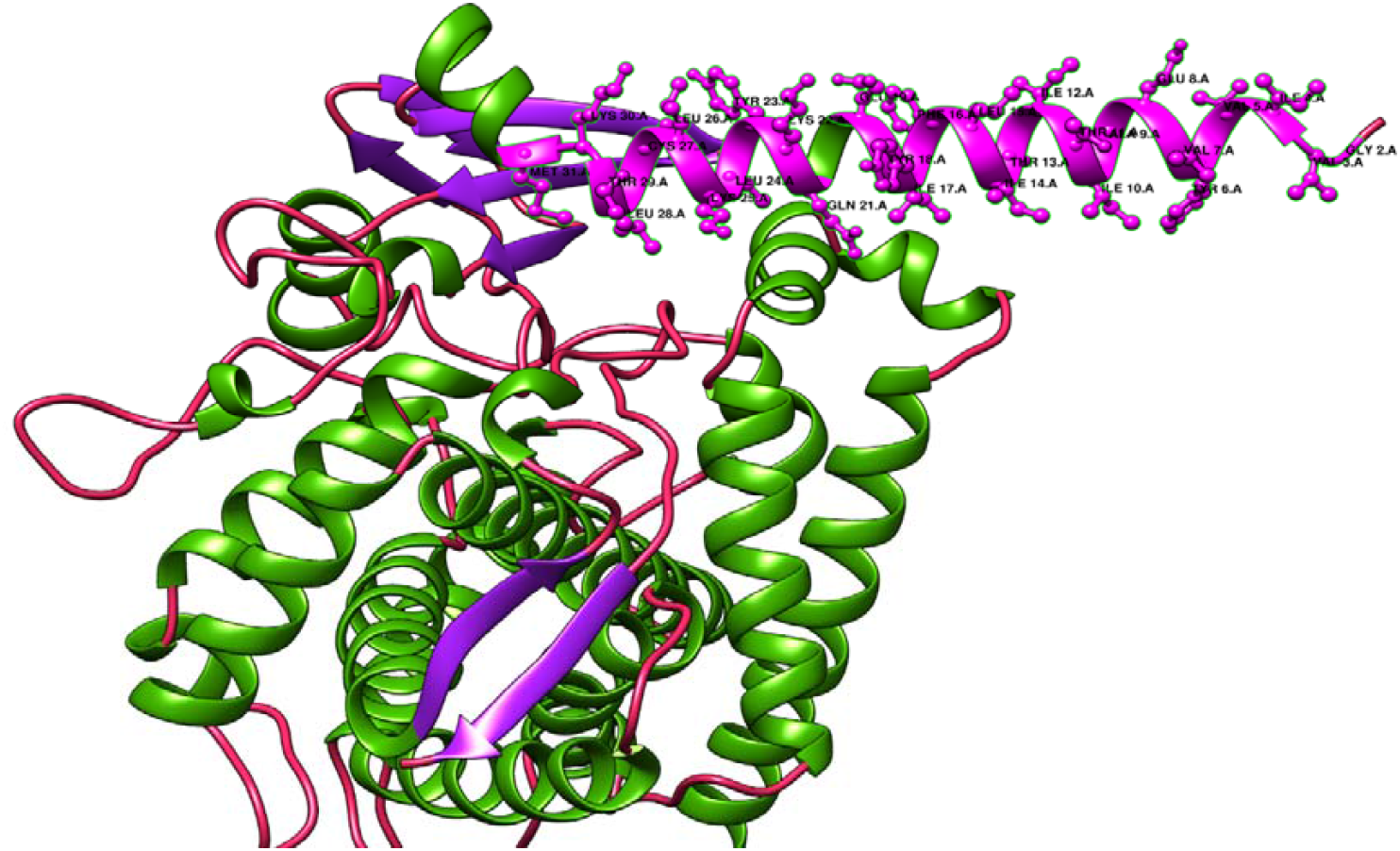
positively selected sites (BEB) represented in the AlphaFold model structure of CYP2C42 in squirrel The positively selected residues (1P-33K) are colored in magenta.

The UGT1A family shows more duplication events in primates (human and bushbaby), few in rodents (mouse and squirrel), and elephants, while pseudogenized or lost for most domesticated species, like cats, goats and chickens (no traces of pseudogenes were found by tBlastn).

The CYP3A and CYP2D family underwent huge duplication events across rodents and ungulates, generating 9 paralogs of each in mice. Two pseudogenes are found by tBlastn in humans (CYP2D7 and CYP2D8P) but already proved in literature^12^. The CYP2B6 family is absent in mice, rats and other rodents, although the syntenic region is rich in other CYP families, like the CYP2A, CYP2B, CYP2F, CYP2S, CYP2G in these species.

While the CYP2C family is absent in most species, with pseudogenes detected by tBlastn in cat, pig, cow and horse, and only present in few rodents like mice, rats and squirrels, few primates like human and gorilla, and some Artiodactyla (like the Siberian musk deer and the Yarkand deer).

The GSTP1 genes are absent in most carnivores and ungulates, and present in primates and rodents. A pseudogene was only identified in the cow, and no other domesticated species.

The NAT1 genes are present throughout all the domesticated species, except the dog, but we could not find any trace of pseudogene. The canids in general lack the NAT activity^13^, although it is conserved in most other species (one pseudogene in human found in literature^14^).

## POSITIVE SELECTION

The CODEML analysis revealed great positive selection within the CYP families, the UGT families and the NAT1 family, while there was no positive selection detected in the GSTP1 families, suggesting it may be subject to stronger purifying selection or functional constraint.

In particular, the site model revealed strong positive selection, among the CYP2D6, CYP3A, CYP2B, UGT1A and UGT2B families.

For CYP2D6, comparisons between models M1a vs M2a and M7 vs M8 showed highly significant LRTs (31.47, 36.54 respectively) at sites 81S, 138A, 142L, 320S, and 326P. When analyzing mouse paralogs separately, additional selected sites emerged (67V, 141G,145 S 323R, 329C), suggesting possible sub-functionalization in the mouse lineage, likely associated with the extensive gene duplications observed in rodents. For CYP2B, comparisons between models M1a vs M2a were not statistically significant, while M7 vs M8 model revealed one positively selected site (146S). For the CYP3A family, The M7 vs M8 comparison (LRT=32.49) detected selection at sites 107P, 224V, 239S, and 320E, while M1a vs M2a only detected it at 107P, probably due to the high sensitivity of the M8 model in detecting selection.

For the UGT families, UGT1A showed weaker evidence of selection (sites 90R, 108N detected by the M8 model), while UGT2B had only one common site between the models M2a and M8 (118F) with a moderately significant LRT (12,94 and 15,71 respectively). This suggests more constrained evolution in these genes compared to CYP families.

Meanwhile the branch-site model further revealed strong positive selection, among the CYP2D, CYP2B, CYP2C, UGT1A and NAT1 families, but in specific lineages and genes.

For the CYP2D, it was detected in mouse paralogs with a strong LRT of 39.08 in five sites (138A, 142L, 316H, 320S, 326P), which can suggest rapid evolution of these paralogs, possibly for specialized detoxification functions in rodents.

For the CYP2B, it was detected in goat with the most significant LRT of 80.43 and ω=999 in sites ranging from 24K to 33D and 56K to 58V, and also in CYP2B4 of rabbit, which can reflect an adaptation to plant secondary metabolites in diet.

For the CYP2C, positive selection was detected in Squirrel with LRT=60.33 and ω=999 in the N-terminal region (1P-33K), probably associated with structural adaptations, as well as in rabbit with an LRT=6,37.

For UGT1A, lineage-specific selection was observed in bushbaby (LRT=32.59, ω=30.43) at sites 72V and 288G to 294P, and also in humans (LRT=4,04, ω=24.73) at a single site 171F, which can be related to metabolic specializations in primate species.

For NAT1, it was detected in humans affecting 2 sites (276 E and 195 E) with LRT of 8.82 and ω=96.3, confirming previous studies^15^.

**Table 1:**
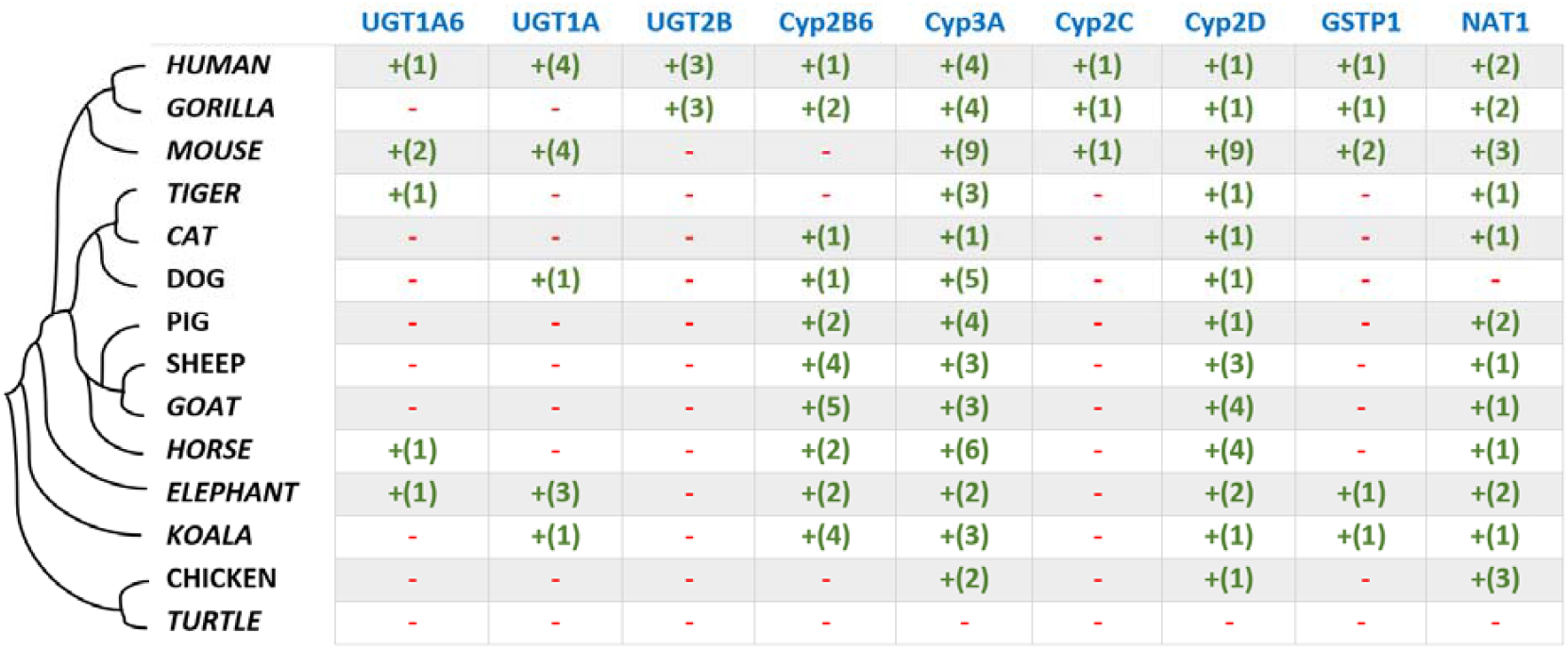
a summary of genes’ duplication and loss across the UGTs, CYPs, GSTP1 and NAT1 +() :present gene family + the number of genes with a paralogous relationship - :lost or pseudogenized

|  | UGT1A6 | UGT1A | UGT2B | Cyp2B6 | Cyp3A | Cyp2C | Cyp2D | GSTP1 | NAT1 |
| --- | --- | --- | --- | --- | --- | --- | --- | --- | --- |
| **HUMAN** | +(1) | +(4) | +(3) | +(1) | +(4) | +(1) | +(1) | +(1) | +(2) |
| **GORILLA** | - | - | +(3) | +(2) | +(4) | +(1) | +(1) | +(1) | +(2) |
| **MOUSE** | +(2) | +(4) | - | - | +(9) | +(1) | +(9) | +(2) | +(3) |
| **TIGER** | +(1) | - | - | - | +(3) | - | +(1) | - | +(1) |
| **CAT** | - | - | - | +(1) | +(1) | - | +(1) | - | +(1) |
| **DOG** | - | +(1) | - | +(1) | +(5) | - | +(1) | - | - |
| **PIG** | - | - | - | +(2) | +(4) | - | +(1) | - | +(2) |
| **SHEEP** | - | - | - | +(4) | +(3) | - | +(3) | - | +(1) |
| **GOAT** | - | - | - | +(5) | +(3) | - | +(4) | - | +(1) |
| **HORSE** | +(1) | - | - | +(2) | +(6) | - | +(4) | - | +(1) |
| **ELEPHANT** | +(1) | +(3) | - | +(2) | +(2) | - | +(2) | +(1) | +(2) |
| **KOALA** | - | +(1) | - | +(4) | +(3) | - | +(1) | +(1) | +(1) |
| **CHICKEN** | - | - | - | - | +(2) | - | +(1) | - | +(3) |
| **TURTLE** | - | - | - | - | - | - | - | - | - |

**Table 2:**
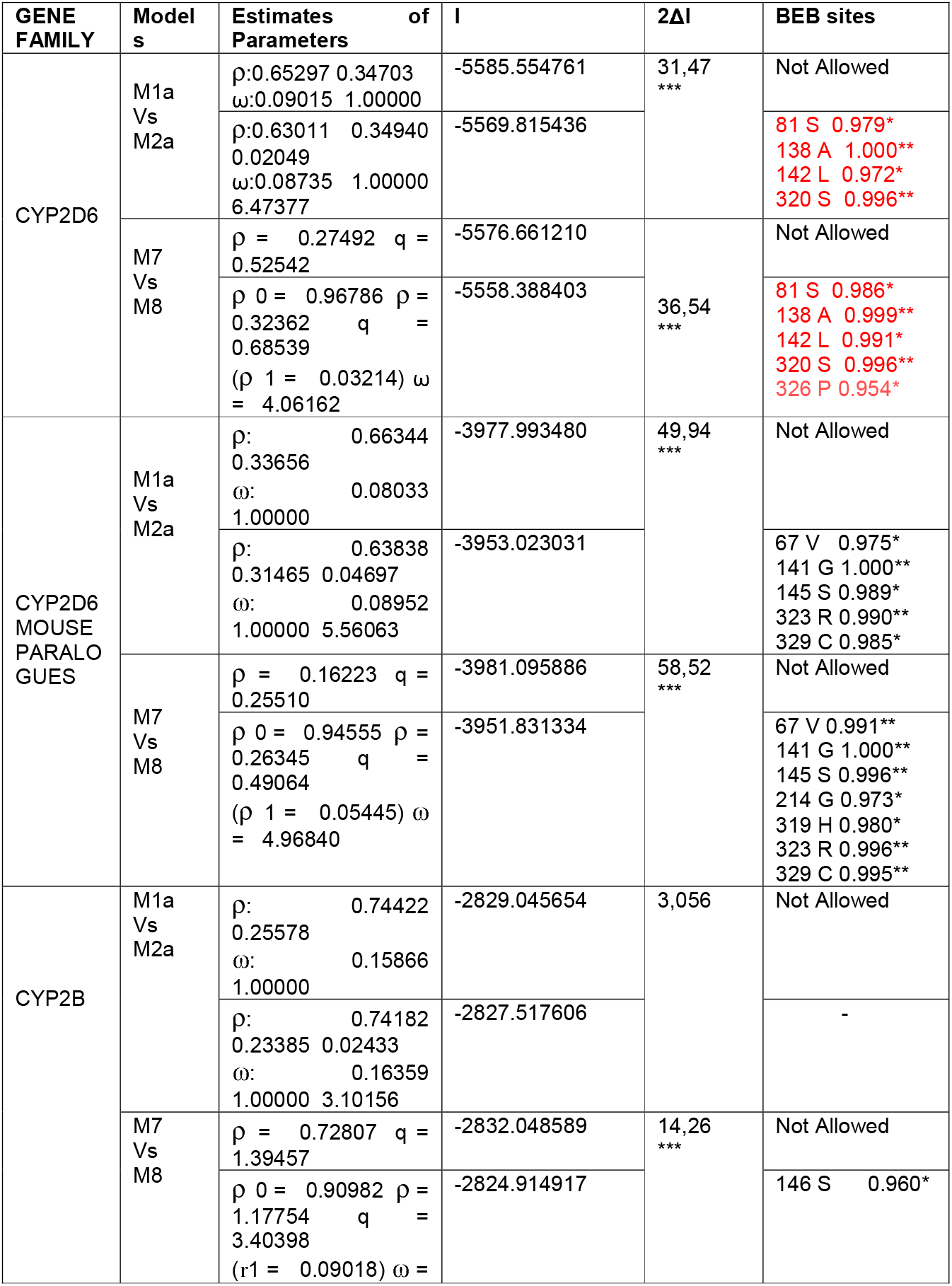
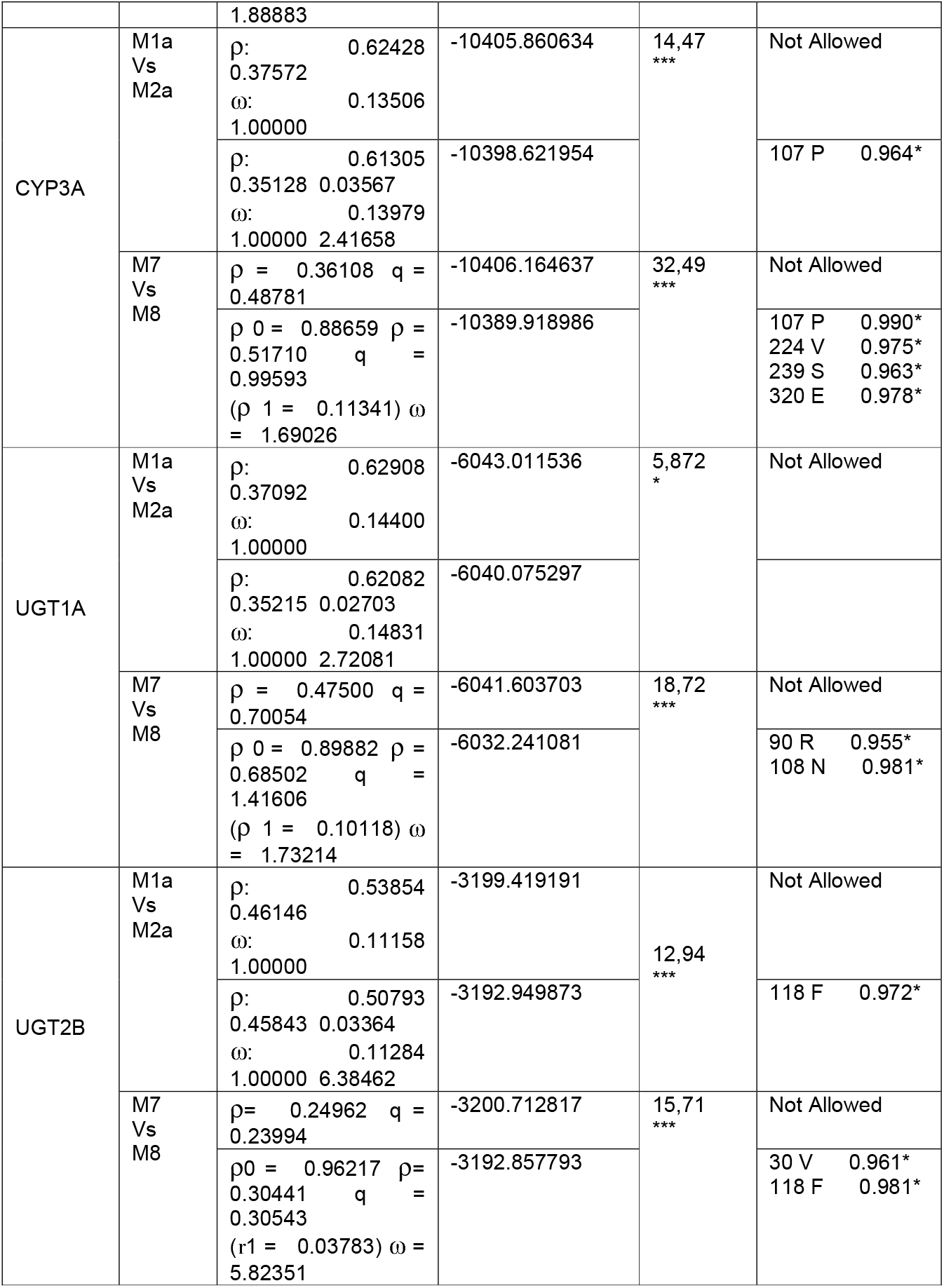
Parameter estimates and likelihood scores for site models for paralogs and orthologs combined l : Log- likelihood values. 2Δl : likelihood ratio test ρ, ρ0, ρ1 : the proportions of codons subject to purifying selection, neutral evolution, and positive selection, respectively. ω : dN/dS for each class (purifying, neutral and positive selection, respectively). *** : significant at p< 0.001 ** : significant at p< 0.01 * : significant at p< 0.05

**Table 3:** Parameter estimates and likelihood scores for branch-site models for paralogs and orthologs combined l : Log- likelihood values. 2Δl : likelihood ratio test ω : dN/dS for each class (purifying, neutral and positive selection, respectively). *** : significant at p< 0.001 ** : significant at p< 0.01 * : significant at p< 0.05

| GENE FAMILY | Foreground branch | Model | Site class | Proportion | Background $\omega$ | Foreground $\omega$ | $l$ | $2\Delta l$ | Positively selected sites |
| --- | --- | --- | --- | --- | --- | --- | --- | --- | --- |

Table3 : Parameter estimates and likelihood scores for branch-site models for paralogs and orthologs combined
|  |  |  |  |  |  |  |  |  | (BEB) |
| --- | --- | --- | --- | --- | --- | --- | --- | --- | --- |
| CYP2D | All paralogs of mouse | Null | 0<br>1<br>2a<br>2b | 0.60375<br>0.26428<br>0.09179<br>0.04018 | 0.07305<br>1.00000<br>0.07305<br>1.00000 | 0.07305<br>1.0000<br>1.00000<br>1.0000 | -<br>5579.<br>4869<br>38 | 39,08<br>*** | Not Allowed |
|  |  | Alternative | 0<br>1<br>2a<br>2b | 0.62929<br>0.33456<br>0.02360<br>0.01255 | 0.08989<br>1.00000<br>0.08989<br>1.00000 | 0.08989<br>1.00000<br>9.07594<br>9.07594 | -<br>5559.<br>9432<br>36 |  | 138 A 1.000**<br>142 L 0.952*<br>316 H 0.972*<br>320 S 0.998**<br>326 P 0.962* |
| CYP2B | ENSCHIG00<br>000023417<br>(ens37)<br>/<br>Goat | Null | 0<br>1<br>2a<br>2b | 0.38081<br>0.13645<br>0.35540<br>0.12734 | 0.12990<br>1.00000<br>0.12990<br>1.00000 | 0.12990<br>1.00000<br>1.00000<br>1.00000 | -<br>2817.<br>2092<br>85 | 80,43<br>*** | Not Allowed |
|  |  | Alternative | 0<br>1<br>2a<br>2b | 0.65841<br>0.25244<br>0.06444<br>0.02471 | 0.12964<br>1.00000<br>0.12964<br>1.00000 | 0.12964<br>1.00000<br>999.0000<br>0<br>999.0000<br>0 | -<br>2776.<br>9904<br>72 |  | 24 K 0.988*<br>25 I 1.000**<br>26 H 1.000**<br>27 E 1.000**<br>28 I 0.988*<br>29 Q 1.000**<br>30 R 0.990*<br>31 Y 0.989*<br>32 S 0.993**<br>33 D 1.000**<br>56 K 0.989*<br>57 W 0.994**<br>58 V 0.999** |
|  | CYP2B4<br>/<br>Rabbit | Null | 0<br>1<br>2a<br>2b | 0.45332<br>0.14362<br>0.30608<br>0.09698 | 0.11720<br>1.00000<br>0.11720<br>1.00000 | 0.11720<br>1.00000<br>1.00000<br>1.00000 | -<br>2812.<br>8859<br>03 | 29,73<br>** |  |
|  |  | Alternative | 0<br>1<br>2a<br>2b | 0.44450<br>0.13983<br>0.31619<br>0.09947 | 0.13189<br>1.00000<br>0.13189<br>1.00000 | 0.13189<br>1.00000<br>50.93030<br>50.93030 | -<br>2798.<br>0162<br>77 |  | 55 V 0.987*<br>68 S 0.960*<br>69 S 0.959*<br>70 S 0.996**<br>74 A 0.971*<br>75 R 0.988* |
| CYP2C | CYP2C42<br>/<br>Squirrel | Null | 0<br>1<br>2a<br>2b | 0.30687<br>0.21057<br>0.28618<br>0.19638 | 0.13359<br>1.00000<br>0.13359<br>1.00000 | 0.13359<br>1.00000<br>1.00000<br>1.00000 | -<br>1544.<br>9000<br>88 | 60.33<br>*** | Not allowed |
|  |  | alternative | 0<br>1<br>2a<br>2b | 0.36468<br>0.26720<br>0.21245<br>0.15566 | 0.13275<br>1.00000<br>0.13275<br>1.00000 | 0.13275<br>1.00000<br>999.0000<br>0<br>999.0000<br>0 | -<br>1514.<br>7363<br>89 |  | 1 P 0.994**<br>2 V 0.977*<br>4 V 0.996**<br>5 L 0.994**<br>6 V 0.999**<br>7 L 0.993**<br>8 T 0.995**<br>9 L 1.000**<br>10 S 1.000**<br>11 S 1.000**<br>12 L 0.999**<br>13 L 0.999**<br>14 L 0.993**<br>15 L 0.995**<br>16 S 0.999**<br>17 L 0.968*<br>18 W 0.999**<br>19 R 0.997**<br>21 G 1.000**<br>22 N 0.993**<br>23 T 0.999**<br>24 L 0.996**<br>25 Q 0.992**<br>26 I 0.999** |
|  |  |  |  |  |  |  |  |  | 27 Y 0.999**<br>28 M 0.993**<br>29 K 0.997**<br>30 D 0.997**<br>31 I 0.965*<br>32 G 1.000**<br>33 K 0.972* |
|  | CYP2C16<br>/<br>Rabbit | Null | 0<br>1<br>2a<br>2b | 0.11942<br>0.17523<br>0.28588<br>0.41948 | 0.08977<br>1.00000<br>0.08977<br>1.00000 | 0.08977<br>1.00000<br>1.00000<br>1.00000 | -<br>1548.<br>3224<br>99 | 6,37<br>* | Not allowed |
|  |  | Alternative | 0<br>1<br>2a<br>2b | 0.26524<br>0.40627<br>0.12975<br>0.19874 | 0.08283<br>1.00000<br>0.08283<br>1.00000 | 0.08283<br>1.00000<br>999.0000<br>0<br>999.0000<br>0 | -<br>1545.<br>1371<br>86 |  | 8 T 0.980*<br>48 K 0.993**<br>85 G 0.992** |
| UGT1A | UGT1A8<br>/<br>Human | Null | 0<br>1<br>2a<br>2b | 0.57450<br>0.34045<br>0.05341<br>0.03165 | 0.14158<br>1.00000<br>0.14158<br>1.00000 | 0.14158<br>1.00000<br>1.00000<br>1.00000 | -<br>6042.<br>3807<br>18 | 4,04<br>* | Not Allowed |
|  |  | alternative | 0<br>1<br>2a<br>2b | 0.62087<br>0.37001<br>0.00572<br>0.00341 | 0.14083<br>1.00000<br>0.14083<br>1.00000 | 0.14083<br>1.00000<br>24.73125<br>24.73125 | -<br>6040.<br>3565<br>64 |  | 171 F 0.952* |
|  | ENS40<br>ENSOGAG0<br>0000028447<br>/<br>Bushbaby<br>(ens40.B) | Null | 0<br>1<br>2a<br>2b | 0.51257<br>0.28987<br>0.12619<br>0.07137 | 0.13531<br>1.00000<br>0.13531<br>1.00000 | 0.13531<br>1.00000<br>1.00000<br>1.00000 | -<br>6038.<br>4523<br>73 | 32,59<br>*** | Not Allowed |
|  |  | alternative | 0<br>1<br>2a<br>2b | 0.59795<br>0.33154<br>0.04536<br>0.02515 | 0.13801<br>1.00000<br>0.13801<br>1.00000 | 0.13801<br>1.00000<br>30.43274<br>30.43274 | -<br>6022.<br>1565<br>91 |  | 72 V 0.993**<br>288 G 0.955*<br>291 P 0.982*<br>292 Q 0.971*<br>293 T 0.996**<br>294 P 0.983* |
| NAT1 | NAT1<br>/<br>Human | Null | 0<br>1<br>2a<br>2b | 0.69186<br>0.30814<br>0.00000<br>0.00000 | 0.14030<br>1.00000<br>0.14030<br>1.00000 | 0.14030<br>1.00000<br>1.00000<br>1.00000 | -<br>4550.<br>6022<br>59 | 8,82<br>* | Not Allowed |
|  |  | alternative | 0<br>1<br>2a<br>2b | 0.00000<br>0.00000<br>0.69834<br>0.30166 | 0.13850<br>1.00000<br>0.13850<br>1.00000 | 0.13850<br>1.00000<br>96.30948<br>96.30948 | -<br>4546.<br>1923<br>32 |  | 276 E 0.976*<br>195 E 0.977* |

## VISUALISATION OF CYP2D6 POSITIVELY SELECTED SITES

The mapping of positively selected sites onto the crystal structure of human CYP2D6 (PDB: 3TDA) revealed intriguing insights into the functional and evolutionary significance of these residues: 81S, 300A, 304S, 138A, 142L. Notably, the fact that two of the identified sites, alanine at position 300 and serine at position 304 are in interaction with the inhibitor Prinomastat (PN0). The serine forms a hydrogen bond (2.97 Å) with PN0, while the alanine participates in a non-contact interaction (3.3 Å), both contributing to the substrate/inhibitor binding^16^. In contrast, the other positively selected sites (81S, 138A, 142L) were located on surface-exposed helices and coils, distant from the active site of the heme group.

While the CYPC42 of squirrel shows positive selection affecting the N-terminal segment (1P-33K), forming a flexible helix that interacts with the endoplasmic reticulum membrane, which is critical for structural stability and membrane anchoring in cytochrome P450 enzymes.

## DISCUSSION

The huge duplication and loss events within the CYP, UGT, GSTP1, and NAT1 gene families reflect the evolutionary flexibility of these enzymes between species. For example, the CYP2D and CYP3A families had massive duplication events in rodents and ungulates, probably reflecting the adaptive response of these species to the high diversity of xenobiotics encountered in their environments, which may have led to the expansion of metabolizing enzymes through gene duplication followed by sub- functionalization. In contrast, the UGT1A and UGT2B families showed extensive gene loss and pseudogenization in carnivores and ungulates. It was already demonstrated in earlier studies that Cats lack a huge pattern of the UGT1A isoforms compared to humans^17^, this being one of the reasons they have very limited capacity to metabolize the acetaminophen(paracetamol), since This latter is mainly eliminated through glucuronidation by the UGTs^18^. Similarly, dogs exhibit limited capacity for certain acetylation reactions due to differences in NAT expression compared to humans^13^. These trends indicate that the evolutionary histories of these enzymes are dictated both by environmental forces and species-specific metabolic requirements.

The presence of pseudogenes in the conserved syntenic regions also supports the suggestion that these families of genes have undergone frequent functional differentiation. For example, the replacement of UGT2B paralogues by UGT2A genes in goats and cattle reflects a functional alteration maybe due to dietary or environmental stresses. This may represent an event of gene loss compensated by functional equivalents, or with different substrate specificities.

The difference between the site model and branch-site model results illustrates the complexity of evolutionary processes and the difficulty of interpretation of results obtained thanks to the tools to study them. While the site model broadly identified conserved positively selected sites, the branch-site model revealed lineage-specific adaptations not revealed in the general analysis. This suggests that the two models are complementary to one another, that the site model provides an overall picture of selection pressures, and that the branch-site model provides finer details of lineage-specific evolution. Future studies would be strengthened by employing multiple models to account for the full spectrum of evolutionary dynamics.

The identification of 2 positively selected sites Ala-300 and Ser-304 within the binding site of CYP2D6 is in line with the previously known function of this protein to metabolize most xenobiotics, where minor alterations in the binding-site residues would significantly change substrate specificity or catalytic activity among species. These helix I residues interact via a backbone hydrogen bond, with Ala300’s proximity to Asp301 (which binds cationic substrates like thioridazine) helping to maintain the active site geometry while allowing the structural flexibility needed to accommodate diverse ligands, as shown by helix I’s conformational changes in different ligand-bound states, as seen in the outward bulging of helix I in closed states like in the prinomastat complex^11^. Species-specific variations at these sites may explain differences in drug metabolism, as for example humans vs. mice, supporting the hypothesis that rapid evolution of CYP2D6 reflects specific dietary or xenobiotic challenges. For instance, Wild-type mice do not metabolize debrisoquine through 4-hydroxylation, while transgenic mice expressing a humanized CYP2D6 are able to perform this metabolic reaction^19^.

Compared to CYP2D6, where selected sites include active-site residues (Ala300, Ser304), the squirrel’s CYP2C42 shows selection in non-catalytic region, that is often involved in membrane anchoring and substrate access, suggesting divergent evolutionary pressures, possibly affecting structural or regulatory adaptations.

The spatial distribution of the squirrel’s CYP2C42 helix and the 3 other positively selected sites (81S, 138A, 142L) of human’s CYP2D6 is consistent with patterns observed in other rapidly evolving proteins, where positive selection often targets peripheral regions involved in protein-protein interactions or structural stability^20^. For example, surface residues may influence enzyme stability in different physiological environments, like temperature and pH, or mediate interactions with redox partners like cytochrome P450 reductase. The absence of direct involvement in catalysis suggests these sites could underlie lineage-specific adaptations to environmental or dietary pressures without disrupting core enzymatic function.

This study opens many new areas of investigations. Most notably, functional assays need to test experimentally the effect of the positively selected sites on enzyme activity and substrate recognition. Site-directed mutations could be employed to determine their precise functional effects. Secondly, widening the scope of analysis to cover additional species and additional gene families would give an even better understanding of the evolutionary dynamics of such enzymes. Moreover, in human medicine, there is a big difference in the ability of individuals to metabolize a drug. Carrying out the same study on a large scale on several hundred to thousands of human genomes (and other animal species such as dogs or cattle) would be an avenue for the future. In particular, incorporating intra-species variation (for example in term of copy number variation and structure comparisons, like superimposing protein models from various species, would give additional data about how these positively selected residues influence protein structure and function. Lastly, matching genomic information with ecological and dietary information would determine the specific environmental and metabolic selective pressures that may have caused the observed patterns of adaptive evolution.

## CONCLUSION

This study highlights the very diverse evolutionary processes and positive selection have in shaping the species-specific drug-metabolizing enzyme diversity. Through gene families CYP, UGT, GSTP1, and NAT1, we uncovered the common gene duplication, loss, and pseudogenization events, reflecting adaptive responses to dietary and environmental pressures. The identification of sites that are positively selected in critical enzymes like CYP2D6, particularly those involved in substrate binding and structural stability, underscores the functional importance of these evolutionary changes.

